# Estimating animal density with camera traps: a practitioner’s guide of the REST model

**DOI:** 10.1101/2021.05.18.444583

**Authors:** Yoshihiro Nakashima, Gota Yajima, Shun Hongo

**Affiliations:** College of Bioresource Science, Nihon University, 1866 Kameino, Fujisawa, Kanagawa 252□0880, Japan; Graduate School of Asian and African Area Studies, Kyoto University, Kyoto 606□8501, Japan

**Author notes:** Corresponding author: E□mail.

## Abstract

Camera traps are increasingly popular in wildlife research and have the potential to be reliable and cost-effective for estimating animal density. Although density estimation using this automatic technique has long been restricted to species with individually recognizable markings, several analytical approaches have been proposed to target animals lacking such markings (i.e., unmarked populations). Among these approaches, the random encounter and staying time (REST) model may be an efficient and cost-effective approach, even though the procedures for the implementations have not yet been shared with researchers. This paper presents a working protocol for implementing the REST model our research group has been developing. We also present the R code to perform parameter estimation with a maximum likelihood and Bayesian approach. We suggest that this model has potential for further development. We strongly hope that this paper will encourage many researchers to use the REST for density estimation in a wide variety of species in various habitats and make a significant contribution to advancing wildlife conservation and management.

## Introduction

Reliable and accurate information on animal population density and its spatial and temporal variations provide a basis for elucidating the patterns and processes of animal populations and community dynamics (Thompson et al. 1998). Thus, such information is fundamental to ecological studies and is critical for the proper conservation and management of wildlife and habitats (Nichols and Williams 2006). Accordingly, researchers have tried to establish an efficient, cost-effective, and reliable approach to estimate animal density for many years. Nonetheless, currently available methods, most of which rely on visual detection of animals or animal signs, are often very costly (e.g., direct counts) or are not always reliable (e.g., dung counts) (Plumptre 2000). Therefore, further efforts are necessary to establish new alternatives using innovative field and analytical techniques to mitigate tradeoffs between survey efficiency and reliability.

In recent years, commercially available camera traps with infrared sensors have become more efficient and less expensive and have become increasingly popular in wildlife research (Burton et al. 2015). Although camera traps can be used for numerous objectives and have led to many findings (McCallum 2013), they also open up new opportunities in estimating the density of ground dwelling mammals. In the past, density estimation using this tool has been limited to recognizing animal individuals and applying spatially explicit capture-recapture models (Royle et al. 2009, Borchers et al. 2014). In recent years, however, several analytical approaches applicable to unmarked populations (i.e., species lacking individually recognizable markings) have been developed, making it possible to target a wide range of species (Gilbert et al. 2020). Rowcliffe et al. (2008) pioneered this work and presented a random encounter model (REM). This model formulates the relationship between animal density, the area of camera detection zones, trapping rates (the number of animal detections per unit time), and animal movement speed based on an ideal gas model. Subsequently, Chandler and Royle (2013) presented a spatial count (SC) model to estimate the density of unmarked populations based on the spatial pattern of animal detection. More recently, several novel approaches have been proposed one after another (Ramsey et al. 2015, Howe et al. 2017, Campos□Candela et al. 2018, Moeller et al. 2018, Nakashima et al. 2018, Luo et al. 2020), establishing one of the most active areas of camera trap research (Gilbert et al. 2020).

The random encounter and staying time (REST) model proposed by Nakashima et al. (2018) is one approach to estimating animal density exclusively utilizing camera trapping data without individual animal recognition. The REST model uses staying times in a predefined focal area within a camera detection zone to account for animal movement speed, which can be easily measured using camera trapping data. This approach may provide a good balance between the efficiency of the survey and the reliability of the estimates. Palencia et al. (2021) compared three density estimation models, the REM, CM DS (distance sampling with camera traps, Howe et al. 2017), and REST, and confirmed that these provided consistent results in most situations. They also suggest that the REST model may be more cost-effective than the other models. However, it may be unsuitable for animals with low density. Nonetheless, this model is still new, and the necessary procedures to acquire the required data have not yet been fully established. If a detailed working protocol is shared among practitioners, density estimation using this approach will be more effective and promising.

In this paper, we concisely summarize the theory of the REST model and the working protocol that our research group has been developing. We also present the R code necessary for maximum likelihood estimation and Bayesian approaches. Our work focuses on how to accomplish data collections and estimate density efficiently. Refer to Nakashima et al. (2018) for the statistical details of the model.

### Model framework and the assumptions

The REST model is based on the formula as follows.

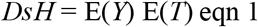

where *D* is animal density, *s* is the size of the area that a camera trap monitors (hereafter focal area), *H* is the effective research period, *E*(*Y*) is the expected number of animals passing through the focal area, and *E*(*T*) is the expected time an animal stays within the focal area (hereafter staying time).

This formula is derived as follows: Let us assume that one camera trap with a focal area *s* is set at a random location within a 2□dimensional habitat with animal density *D*. The number of animals in the focal area at any given time can be expected to be *D* × *s*, by definition of density. The total length of time the animals stay within the focal area during the effective research period *H* is then *D* × *s* × *H*. On the other hand, if the camera trap is designated to take video, one can estimate the expected number of animals passing through the focal area *E*(*Y*) and the expected staying time within the focal area *E*(*T*). In this case, the total length of the animals staying within the focal area during the entire research period was *E*(*Y*) × *E*(*T*). Thus, we obtained Equation 1. For more details, refer to Nakashima et al. (2018).

The REST model adheres to the following seven assumptions: (1) camera traps are randomly placed, (2) when the animal passes through a predefined focal area, camera traps must certainly detect the animals without delay throughout the research period, (3) animal density must remain constant during the research period, (4) animal movement and behavior must not be affected by camera traps, and (5) the data must be independent. Furthermore, when density is estimated using a likelihood-based approach (see below), (6) the number of animal passes and staying times within the focal area must follow a parametric probability distribution. In practice, it is also necessary that (7) the effective research period can be estimated (see below).

In the following, we will follow the series of processes up to density estimation in chronological order and describe practical approaches to satisfy these assumptions and the procedures for efficiently carrying out the work in the real world.

### Considerations before surveys

Before the survey, we recommend considering whether absolute animal density (the number of individuals per unit area) is essential to achieve the research objective. In some situations, it may be sufficient to obtain information on relative animal density (i.e., an index proportional to the absolute density) in actual conservation and management. For example, various statistical models have been used to estimate occupancy or abundance using various approaches (Iijima 2020), including camera trapping. Although these approaches may rely on relatively strong assumptions regarding an animal’s ecological state or detection processes (Nakashima 2020), the research cost required by these models is much lower than the estimated absolute density (Jones 2011).

Suppose that density estimation is necessary for these objectives. In this case, the next step is to carefully consider whether the REST model may be the best choice for estimating the density of the target animals. If the target animal is individually recognizable, even partially, a more accurate and precise method may be available (e.g., spatial mark–resight (SMR) model, Chandler and Royle 2013, Sollmann et al. 2013). Even if individual recognition is impossible, other models may be more suitable. For example, if camera traps rarely detect target species due to low density, distance sampling with camera traps (Howe et al. 2017) or the SC model (Chandler and Royle 2013) may be more promising. The REST model only uses records wherein an animal passes through a predefined focal area in the camera trap’s field of view. This restriction can not enhance working efficiency, but it also reduces the sample size, making its application challenging to species with low densities (Palencia et al. 2021).

Another critical aspect is whether the target species meet the assumptions of the model. In particular, one should pay special attention if (7) the effective research period can be estimated. Here, the effective research period refers to the length of time a camera can potentially detect animals.

To clarify this, we consider the case of targeting a semi□arboreal species. When a semi□arboreal species is in a tree, the probability of the camera detecting the animal is zero, which is substantially the same as if the cameras did not function for that period. Thus, it is necessary to exclude the arboreal period of these animals from the total research period to determine the actual research period. The REST model can only be applied to species if we can estimate the duration of the arboreal period. Note that even fully ground dwelling mammals are not always detectable by camera traps in the real world. Consider that deer are entirely diurnal, staying in one place to sleep at night. In this case, even though the density of deer does not vary between day and night, very few videos were captured at night (we all know this experientially). Thus, as in the case of semi□arboreal animals, one cannot include the inactivity period of ground dwelling animals in the effective research period.

How can one estimate the proportion of time that animals are active (or inactive)? Rowcliffe et al. (2014) showed that the capture rate of animals during a given period within a day is proportional to the number of active individuals. Thus, if we can assume that all population members are active at a given time within a day, it is possible to estimate the activity proportion from the camera trapping rate (see Rowcliffe et al. 2014 for more details). However, assumption (7) may limit the range of species that the REST model can be applied to, although this restriction is the same in CM DS and REM.

### Preparation for camera traps

Once the REST model has been chosen to perform the density estimation, one needs to purchase a camera trap model suitable for the REST model or check if the owned camera models are sufficiently efficient for the applications.

At a minimum, the camera traps used should have the following features: First, the model must have a video recording function or capture still images within a short time interval. Second, the model has a minimal time lag between the sensor response and video shooting to detect passing animals. Third, it is better to check the camera’s field of view in real-time to ensure that the camera trap properly monitors the predetermined focal area. Fourth, since the REST model requires a relatively large number of sampling locations, the price per unit should be low enough to allow the purchase of a sufficient number of cameras on a budget. Unfortunately, not many commercially available cameras meet all of these requirements. To the best of our knowledge, the most suitable camera may be Browning’s camera (e.g., browning strike force, https://browningtrailcameras.com/). This camera satisfies all of the abovementioned requirements. In the future, we hope to see camera trap models designed specifically for the REST model.

The number of camera traps required is also an essential aspect, although it may depend on the target species and habitats. In general, it is costly to obtain sufficient data to estimate the expected number of animal passes (Schaus et al. 2020). While data on staying times and activity proportion are available for each animal pass, the number of animal passes can only be obtained from the number of camera locations. Additionally, the number of animal passes may be highly variable among camera locations, reflecting a higher heterogeneity of animal space use. Therefore, keep in mind that the key to successful applications is securing a sufficient number of camera locations. If a sufficient number of camera traps are not available, replacing the cameras during the research period (although it requires considerable labor cost) may be an effective way to survey with a limited number of cameras.

According to the Monte Carlo simulations by Nakashima et al. (2018), data should be obtained from at least 25 camera locations, preferably 50–100. The CV should be less than 10% for proper wildlife management (Williams et al. 2002). To achieve this, installing camera traps at more than 100 locations may be necessary (Bessone et al. 2020). A large amount of survey effort may be necessary to obtain accurate and precise information, particularly for low-density species (Cappelle et al. 2021).

### Determining the survey area and the survey design

In the next step, the survey area and design of camera placements should be determined. The optimal research design depends on the situation, but the following principles may help determine the survey design.

When targeting more homogeneous (often small) habitats, one may conduct complete random or systematic sampling from the entire study area using GIS software. The REST model assumes that the data are independent (Assumption 2). It is better to maintain a certain distance between the cameras to meet this assumption. Completely random sampling requires an arbitrary setting of a minimum distance between cameras. In addition, there is often some bias in the characteristics of the study area even though it appears to be homogeneous (e.g., the soil may be more fertile in the south). Thus, systematic sampling (e.g., setting cameras at equal intervals to cover the entire survey area) may often be less problematic than random sampling. The “two□stage sampling method” may also be effective. This method is conducted by dividing the survey area into meshs in advance and then performing random sampling from each mesh (Nakashima et al. 2013).

If the study area exhibits higher heterogeneity, the survey should be designed according to the characteristics of the habitat. For example, suppose the study area is a mixture of two habitat types, one option is calculating each area in advance and allocating camera locations according to their proportions (i.e., the proportional allocation for stratified sampling). Then, the mean density over the entire study area can be calculated from the density estimates in each layer and the proportion of the area. The problem with this method is that estimating population variance (estimating the precision) may be complicated, but if one uses Bayesian estimation for the parameters, no substantial problems are likely to arise. If some habitat characteristics may affect animal density (or animal staying time), consider the effect of the habitats by incorporating covariates into the model (Nakashima et al. 2020). In this case, extrapolating the relationship between habitat characteristics and density (or staying time) can estimate the number of animals within the entire study area.

### Designations of a focal area

It is also necessary to consider defining a ‘focal area’ where camera traps certainly detect passing animals. The location, size, and shape of the focal area may substantially influence density estimations. Therefore, the focal area should be carefully designated according to the camera model and the characteristics of the target animals.

The detection probability of a camera trap relies on the animal’s position relative to the camera; the closer the distance and the more central the angle of view, the higher the detection probability (see Rowcliffe et al. 2011). In addition, the detection probability more rapidly decrease with the distance from the camera for smaller animals. Therefore, the focal area should be set at the center of the field of view and not too far away. A practical method for the designations may apply the approach of Rowcliffe et al. (2011) and Hofmeester et al. (2017), measuring the distance and angle of animals where the animals are first detected. Assuming that animal space use is homogeneous within the field of view, the area with the maximum number of detections is considered to have the highest detection probability. Therefore, the focal area within the area should be selected.

The shape of the focal area may also affect estimation. As mentioned above, a likelihood-based estimation requires that (6) the number of animal passes and staying times within the focal area must follow a parametric probability distribution. When the animal’s movement speed variance is small, depending on the shape of the focal area (e.g., square), it may be difficult to attribute the staying times to an existing probability distribution. Nonetheless, our simulation results show that large variation in animal movement speed does not significantly impact the estimate. Additionally, our simulation results suggest that an equilateral triangle may be an easy solution to this issue.

When targeting medium-and large sized animals using a camera model used by the authors’ research group (Browning Strike Force Pro), a desirable designation is shown in Figure 1 for medium-and large sized mammals (body mass > 1 kg). This designation is based on the results of prior experiments using domesticated dogs (Yajima and Nakashima 2021). This designation may also be valid for Bushnell cameras, which are the more commonly used camera traps (see Yajima and Nakashima 2021 for details).

**Figure 1.**
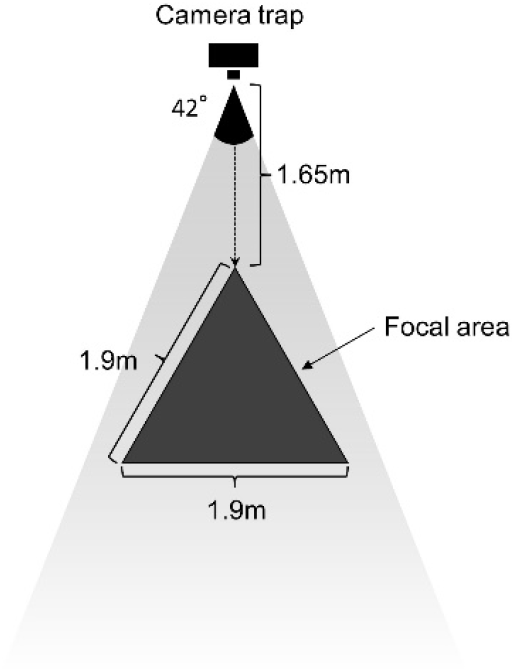
A schematic diagram showing the designation of a focal area where camera traps should certainly detect animals without delay. The focal area is an equilateral triangle with a side of 1.9 m, and the distance from the camera to the nearest vertex is 1.65 m. The camera used (Browning Strike Force Pro) has a sensor detection range of about 42°, and the semi□transparent triangle in the diagram indicates the camera field of view.

### Implementation of survey

Once the survey design is ready, it is time to start a field survey. Take out the camera from the storeroom, insert batteries, and set up camera modes. There are a few configuration points to note. First, camera traps allow the user to set a time interval during which the camera sensor is inactive (the shooting interval). Keep the shooting interval as short as possible to better satisfy certain detection during an effective research period (Assumption 2). Second, reduce censored data, such as data where animals remained in the focal area after filming, to reliably estimate staying times within a focal area by setting up video length; it should not be too short. From our experience, videos longer than 15 s may be more desirable.

During a field survey, one should walk to a planned location with a handy GPS and start camera trap installation without arbitrarily choosing camera positions. In reality, however, the planned location may be inaccessible or unsuitable habitat for camera set-up. It is advisable to determine in advance the principles of how to deal with such situations. For example, “If access is difficult, set up as close as possible,” or “If the planned point is an unsuitable location, move it 30 m to the north.” Such a procedure would violate a random sampling strategy, but flexibility is always required in the field. When conducting surveys, analyzing data, and interpreting the obtained results, these constraints should be considered. Of course, one should follow the principle of random sampling whenever possible. For example, one should not deliberately choose a position where animals are likely to be captured or use baits to attract animals.

Once the camera position is determined, the camera should be mounted so that the predetermined focal area can be successfully captured. We adopted the following installation protocol: (1) Prepare a rope (preferably white) in advance equal to the length of the outline of a predetermined focal area. (2) Use the rope to define the focal area’s outline clearly. (3) Fix the camera firmly while adjusting the height and angle of the camera to maintain the focal area within the angle of view. (4) Film the focal area using the manual shooting function of the camera (the filmed image is referred to as the reference image henceforth). 5. After retrieving the camera traps, the reference image is superimposed on videos of animals to measure whether they pass through the focal area and how long they stay there. A video describing the work has been uploaded to YouTube (https://youtu.be/pUa9rgxSGVA). This method may enhance survey efficiency but is not without its constraints. In particular, if the camera angle changes for some reason (e.g., disturbed by animals) during the survey period, the following data will be unusable. Furthermore, if some animals frequently disturb camera traps, it may be necessary to raise the position at a height that the animals cannot reach. On the one hand, such an installation may make it easier to measure the staying time within the focal area. On the other hand, it may not maximize detection probability since it is more desirable to install a camera parallel to the ground (Yajima and Nakashima 2021). In our experience, it seems easier to achieve a balance when the camera is placed at waist height.

### Data management and measurements

After the planned period has elapsed, retrieve the data and move on to the task in a laboratory.

The first step in the laboratory is to identify the animals captured in the videos. Recently, various applications and software packages have been available to manage camera□trapping data, while a few software packages can handle videos. Currently, we use Timelapse2 (Greenberg, 2016) for species identification. This software has many advantages, including supporting video data, fast data loading, user□adjustable data formats, and export data as a CSV file.

After all the species were identified using this software, we then export the data to a CSV file. Start measuring staying times the following way, although this process is not yet sophisticated. First, play the “reference video,” which shows the focal area with a rope using free software VLC media player (https://www.videolan.org/index.ja.html), and then pause it. Next, playback a video that captures the animals to be measured (hereafter animal video) with VLC in a different window. Then, use the free software TranspWnds (https://code.google.com/archive/p/transpwnds/downloads) to make the animal video semi-transparent and superimpose it on the reference video. As long as the camera angle does not change during the research period, the backgrounds of these videos should overlap exactly. By referring to the location of the rope in the reference video, it should be possible to determine whether the animal has passed through the focal area and measure the staying time of the animals. Finally, manually input the measured values into a CSV file. For more details on measuring the staying times, refer to the video on YouTube (for setting up the computer environments: https://youtu.be/wqEF_up7EGs; for measuring the staying times: https://youtu.be/s-d81K72yWs).

The REST model assumes that (4) the animal’s movement and behavior must not be affected by camera traps. However, animals may react with cameras and change their behavior. One should use such videos to determine the number of animal passes but exclude them from measuring the staying times. In addition, in some videos, video recording starts after the animal enters the focal area. Such videos should be treated as censored data or excluded from the estimation of staying time unless the delay in the detection does not correlate with the animal’s movement speed.

When inputting the staying time into a CSV file, a video may not correspond to a single passage of an animal. For example, an animal may pass through a focal area multiple times in one video, or another animal may pass through the area. In addition, many camera models automatically stop shooting after a preset time, whether the animal remains within the focal area so that same passing may be recorded in multiple videos. In these cases, one should edit the CSV file so that one video line corresponds to one animal pass. For example, if multiple passes in the same video, copy and paste the video information for the number of passes and enter the staying time for each.

On the other hand, if one passing sequence spans multiple videos, the total staying time might be input in the first line and NA in the other lines. Depending on the camera model, it may not be possible to set the “shooting interval” to zero. In this case, if the same animal stays in the focal area for multiple videos, the staying time should be measured by subtracting the time of entering from that of leaving the focal area. Of course, animals may have left and returned to the focal area during the shooting interval, or individuals may have been replaced. However, it is seldom difficult to determine whether or not they are the same individuals in practice.

Although measuring the staying time is the most labor-intensive process in the applications of the REST model, it is not always necessary to measure the staying times in all the videos. Instead, it is better to measure the CV of the staying times to determine the required amount of data.

The goal is to produce data in a format similar to that shown in Figure 2. Censoring indicates that the video was terminated before the animal left the focal area. We recommend preparing a separate CSV file for camera station information (e.g., research period and habitat covariates).

**Figure 2.**
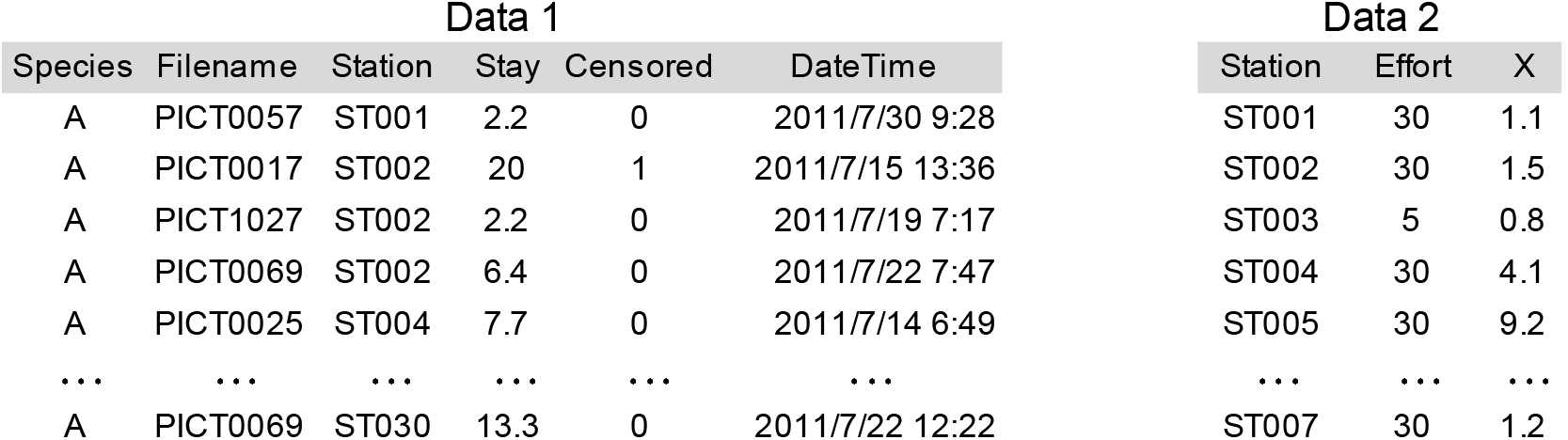
An example of data formats for the REST model. In Data 1, each row of information corresponds to one pass of the animal. Each row contains information about the name of the video file (Filename), the species name (Species), the ID of the camera station (Station), whether it is censored or not (Censored), staying time (Censored = 0), censored time (Censored = 1) (Stay), and the date and time the video was taken (DateTime). On the other hand, Data 2 contains information for each station, where ‘Station’ is the station’s name, ‘Effort’ is the total research period, and ‘X’ is an environment variable for each station.

### Parameter estimations

There are two main approaches to obtaining density estimates, including their variance. One is to calculate the mean density based on Eq. One and perform a nonparametric bootstrapping procedure with delta methods to estimate the variance of each parameter. While this method has the advantage of being intuitive and easy to understand, it underestimates the density when censoring occurs. In addition, the relationship between the environment and density is difficult to estimate. The other is to make inferences based on the likelihood of a model by attributing the number of passes and staying time to a parametric probability distribution. Although this method has a wide range of applications, it requires some knowledge of statistical analysis. In the following, we assume the latter approach in particular.

We discuss some points that need to be considered. As mentioned earlier, camera traps can rarely capture animals when they are inactive. It is necessary to exclude the time that the animals are inactive. However, in high-density animal species, an extremely long stay time is occasionally recorded. Theoretically, we should not use the data when animals are inactive, but in practice, it is often difficult to objectively distinguish whether the data are from active or inactive periods. Whether or not to include such data in estimating the staying time may make a big difference in the density estimates. One realistic approach may exclude outliers by assuming a specific parametric probability distribution (e.g., a log-normal distribution). For example, Nakashima et al. (2018) used the ‘extremevalues’ package (van der Loo 2010) in R to exclude outliers from data. Although it is difficult to justify using a given parametric distribution, the results of the simulations show that this approach may be effective.

All data should be used to estimate the expected number of animal passes and staying times. In many studies, consecutive trapping of the same species at the same station within a given period is often discarded to ensure independent observations (e.g., O’Brien, Kinnaird, & Wibisono, 2003). However, this treatment violates the assumption that camera traps must certainly detect the animals without delay throughout the research period (Assumption 2) and thus should not be performed.

Here is an example of density estimation in R using Data1 and Data 2, as shown above. We will use the ‘tidyverse’ package (Wickham et al. 2019) of R to summarize the data and the ‘bbmle’ package (Bolker and R Development Core Team 2020) to estimate the parameters using a maximum likelihood approach.

First, the data were read in, and the total number of animal passes was counted as follows:

~~~
library(tidyverse)
d1 <- read_csv(“data1.csv”) # Data on staying times
d2 <- read_csv(“data2.csv”) # Information on camera stations
dy <- d1 %>%
   group_by(Station) %>%
   summarize(Y = length(Station)) %>%
   right_join(d2, by = “Station”)
dy$Y[is.na(dy$Y)] <- 0
~~~

Column Y in the data frame dy has information on the number of animal passes at each camera station. We then prepare objects that contain the data required for the REST model:

~~~
stay <- d1$Stay[d1$Censored == 0]   # Staying times (s) not censored
cens <- d1$Stay[d1$Censored == 1]  # Censored
y <- dy$Y # Number of animal passes
area <- 2.67/1000000        #Area of a focal area (square meters)
effort <- dy$Effort*24*60*60 #Total research effort (s) for each camera trap
~~~

To estimate the proportion of active time, we use a point estimate of the approach by Rowcliffe et al. (2014):

~~~
library(“activity”)
library(“lubridate”)
dtime<-as .POSIXlt(d 1 $DateTime)
trad<-2*pi*(hour(dtime)*60*60 + minute(dtime)*60 + second(dtime))/(24*60*60) #
Convert into radian scale
model<-fitact(trad, bw = bwcalc(trad, K = 3), adj = 1.5, reps = 1)
act<-model@act # Activity proportion
~~~

Now, it is time to estimate density. Here, we simply assume that the staying times follow an exponential distribution (with a parameter rate), and the number of passes follows a Poisson distribution (with a parameter lambda). For uncensored staying times, when the exponential distribution has a parameter, ‘mean.stay’, the log□likelihood can be expressed using the R function ‘dexp’ as:

~~~
dexp(stay, rate = 1/mean.stay, log = TRUE).
~~~

The ‘stay’ is a vector of the staying times.

On the other hand, for the data with censoring, since it can be defined as the probability that the unrecorded actual staying time is longer than the censoring time, the log□likelihood is given by using the cumulative density function ‘pexp’ as follows:

~~~
1 □ pexp(cens, rate = 1/mean.stay, log = TRUE))
~~~

The ‘cens’ is a vector of the censored times.

Because the expected value of the number of animal passes is *E*(*Y*) is *D*s*H*/*E*(*T*) from equation 1, its log-likelihood can be written as follows:

~~~
sum(dpois(y, lambda = density*area*effort*act/mean.stay, log = TRUE))
~~~

The ‘y’ is a vector of the number of animals that pass at camera traps (density: Animal density to be estimated; area: the area of the focal area; effort: the total research period; act: the activity proportion).

That is, the total log□likelihood of the model is as follows;

~~~
sum(dexp(stay, rate = 1/mean.stay, log = TRUE)) + sum(1 □ pexp(cens, rate = 1/mean.stay, log = TRUE)) + sum(dpois(y, lambda = density*area*effort*act/mean.stay, log = TRUE))
~~~

Let us try the maximum likelihood estimation of the parameter (minimizing the negative log-likelihood) using the bbmle package in R.

~~~
library(“bbmle”)
minus.log.lik <- function(mean.stay, density){
  -(sum(dexp(stay, rate = 1/mean.stay, log = TRUE)) +
      sum(1 - pexp(cens, rate = 1/mean.stay, log = TRUE)) +
      sum(dpois(y, lambda = density*area*effort*act/mean.stay, log = TRUE)))
}
fit <- mle2(minus.log.lik, start = list(density = 5, mean.stay = 5))
summary(fit)
~~~

To obtain the AIC value for this model, we use the AIC function.

~~~
AIC(fit)
~~~

Assuming that the number of animal passes follows a negative binomial distribution, the function for calculating the minus log-likelihood is given by adding the dispersion parameter alpha in the model as follows:

~~~
library(MASS) # to use the function ‘dnbinom’
minus.log.lik2 <- function(mean.stay, density, alpha){
-(sum(dexp(stay, rate = 1/mean.stay, log = TRUE)) +
     sum(1 - pexp(cens, rate = 1/mean.stay, log = TRUE)) +
     sum(dnbinom(y, mu = density*area*effort*act/mean.stay, size = alpha, log = TRUE)))
}
fit2<-mle2(minus.log.lik2,start = list(density = 5, mean.stay = 5, alpha = 100))
summary(fit2)
AIC(fit2)
~~~

A comparison of the AIC values suggests that the Poisson model may be better than the negative binomial model in this case.

Furthermore, suppose that one wants to determine whether a normalized environmental variable *x*, at camera station *i* may be related to density *D*_1_ (Nakashima et al. 2020). Using an exponential link function ‘exp’, a density at a camera location *i* may be expressed as *D*_1_ = exp(*alpha* + *beta*x_1_*). The values of alpha and beta can be estimated as follows:

~~~
x <- as.numeric(scale(d2$X))
minus.log.lik3 <- function(mean.stay, alpha, beta){
 -sum(dexp(stay, rate = 1/mean.stay, log = TRUE)) - sum(dpois(y, lambda = exp(alpha + beta*x)*area*effort*act/mean.stay, log = TRUE)) #in the case of Negative Binomial distribution
}
fit3<-mle2(minus.log.lik3,start = list(alpha = 5, beta = 1, mean.stay = 5))
summary(fit3)
AIC(fit3)
~~~

The model with the covariates had a larger AIC value, suggesting that animal space use and density did not vary among camera stations. In reality, staying times and density may be influenced by many factors, including random effects. In this case, it may be more helpful to use a Bayesian approach to parameter estimation rather than the maximum likelihood method; We deposited the R code to estimate the parameters and calculate the model selection criterion using JAGS (Plummer 2017) on the Dryad Digital Repository (https://datadryad.org/stash/share/mzsZ4AMe4z8oFPpiwQ0ncnp1updnf8B-ojHUc1rkKF0). Note that specifying censored data differs among software. In particular, the method of giving censored data by JAGS is complicated to understand, so it can be handled with special care.

### Future challenges

In this paper, we presented a practical protocol to apply the REST model to estimate the density of animals without recognizing individuals. The method described here is only an approach at present, and there is room for further improvement. In particular, the measurement of staying time is still in its infancy. It is necessary to develop specialized application software that can efficiently measure the staying time of animals. In the future, the development of automatic measurements using deep learning technology should also be considered. Unfortunately, the authors do not have sufficient capacity and time to develop such procedures at present. If anyone is interested in such an attempt, we would like to contact the authors.

The REST model is a developing approach, and efforts are being made to improve the models’ efficiency, reliability, and applicability. For example, although the REST model depends on certain detection within a focal area, it is necessary to test this assumption in realistic conditions, and if it is not accepted, an approach needs to be established to mitigate the bias arising from bias violations. Moreover, although we use a point estimate of activity proportion based on Rowcliffe et al. (2014), Núñez Antonio et al. (2018) proposed a Bayesian nonparametric approach, which makes it possible to evaluate the uncertainty of the estimations and thus should be incorporated into the estimation procedures. Furthermore, because the REST model has been developed as a likelihood-based model, it can be incorporated into a model that can integrate data from different sources. Indeed, Yokoyama et al. (2020) gathered animal capture data from administration and constructed a state-space model to integrate data from multiple sources to estimate boar density, habitat use, and catchability. Such integrated models would extend the REST model’s further applications and contribute to actual wildlife management. We hope that the REST model will be used in many habitats to facilitate wildlife ecology research and conservation management.

## Supporting information

Data1 required for implementing the REST model

Data2 required for implementing the REST model

R code for the REST model

## Acknowledgments

This study was funded by the Ministry of Education, Culture, Sports, Science, and Technology/JSPS KAKENHI (Grant Numbers JP 15K07487, 16H05661, 18K06430) and the Environment Research and Technology Development Fund (Grant Number 4□1704).

## Notes

https://datadryad.org/stash/share/mzsZ4AMe4z8oFPpiwQ0ncnp1updnf8B-ojHUc1rkKF0

https://www.youtube.com/playlist?list=PLNlLe3RjftYun9Xh2pJOAuisHaTVyEwAB

### Competing Interest Statement

The authors have declared no competing interest.

### Summary of Updates

This version provides a concise and precise description of the entire process from study design to data analysis. It is strongly recommended that readers follow this version of the protocol.

