## Supplementary material for "Estimating animal density with camera traps: a practitioner’s guide of the REST model": R code for the REST model: REST_bayes.R

```
# Estimations with jags -----------------------------------------------------
setwd("D:/Dropbox/00_Project/MSC_manual_REST") # Set your working directory

library("tidyverse")
library("activity")
library("lubridate")
library("jagsUI")
library("loo")

# Setting data -----------------------------------------------------
d1 <- read_csv("data1.csv")
d2 <- read_csv("data2.csv")

dy <- d1 %>% 
  group_by(Station) %>% 
  summarize(Y = length(Station)) %>% 
  right_join(d2, by = "Station")
dy$Y[is.na(dy$Y)] <- 0

# Estimating activity proportion
dtime<-as.POSIXlt(d1$DateTime)
trad<-2*pi*(hour(dtime)*60*60 + minute(dtime)*60 + second(dtime))/(24*60*60)
model<-fitact(trad, bw = bwcalc(trad, K = 3), adj = 1.5, reps = 1)
plot(model)
act<-model@act

# If you exclude the outliers from staying times, perform as follows
#library("extremevalues")
#K <- getOutliers(stay, dist = "lognormal")
#stay <- stay[-which(stay > K$limit[2])]
#cens <- cens[-which(stay > K$limit[2])]

y <- dy$Y
area <- 2.67/1000000          #Area of a camera detection zone
effort <- dy$Effort*24*60*60    #Research effort (sec.) per camera trap (50 days)

# Caution is required to use censored data in JAGS!!!!!!!!!!
is.censored <- d1$Censored # censored (1) or not (0)
stay <- d1$Stay # Staying time
cens <- ifelse(is.censored == 1, stay + 1, stay)

Nstay<-length(stay)
Nstation<-length(y)
datalist<-list(y = y, area = area, effort = effort, is.censored = is.censored, stay = stay, cens = cens, Nstay = Nstay, Nstation = Nstation, act = act)

# Estimation -----------------------------------------------------
# Poisson-exponential model
model.file<-"~/model1.txt"
sink(model.file)
cat("model
      {
      for(i in 1:Nstay){
        is.censored[i] ~ dinterval(stay[i], cens[i])
        stay[i] ~ dexp(rate)
      }
      for(i in 1:Nstation){
        y[i] ~ dpois(lambda[i])
        log(lambda[i]) <- log(area) + log(effort[i]) + log(density) + log(rate) + log(act)
      }
      density ~ dgamma(0.01, 0.01)
      rate ~ dgamma(0.01, 0.01)
      
      # to calculate the WAIC value
      for(i in 1:Nstay){
        loglik.stay[i]<-log(ifelse(is.censored[i] == 1, 1 - pexp(stay[i], rate), dexp(stay[i], rate)))
      }
      for(i in 1:Nstation){
        loglik.y[i] <- log(dpois(y[i],lambda[i]))
      }
      
      }
      ")
sink()
inits<-function(){
  list(density = 5)
}
parameters <- c("density","rate", "lambda", "loglik.y", "loglik.stay")
n.chains <- 3; n.iter<-2000; n.burnin<-1000; n.thin<-2
n.mcmc <- (n.iter - n.burnin)*n.chains/n.thin

model_pois <- jags(datalist, inits,parameters, model.file,
                   n.chains=n.chains, n.iter=n.iter, n.burnin=n.burnin, n.thin=n.thin)
model_pois$summary[1:3,]

# Caluculate the WAIC value
waic(cbind(model_pois$sims.list$loglik.y, model_pois$sims.list$loglik.stay))

# Negative binomial-exponential model (implemented as Poisson-Gamma mixed model)

model.file<-"~/model2.txt"
sink(model.file)
cat("model
      {
      for(i in 1:Nstay){
        is.censored[i] ~ dinterval(stay[i], cens[i])
        stay[i] ~ dexp(rate)
      }
      for(i in 1:Nstation){
        y[i] ~ dpois(lambda[i])
        lambda[i] <- mu[i]*u[i]
        u[i] ~ dgamma(alpha, alpha)
        log(mu[i]) <- log(area) + log(effort[i]) + log(density) + log(rate) + log(act)
        p[i] <- alpha/(alpha + mu[i])
      }
      density ~ dgamma(0.01, 0.01)
      rate ~ dgamma(0.01, 0.01)
      alpha ~ dgamma(0.01, 0.01)
      
      # to calculate the WAIC value
      for(i in 1:Nstay){
        loglik.stay[i]<-log(ifelse(is.censored[i] == 1, 1 - pexp(stay[i], rate), dexp(stay[i], rate)))
      }
      for(i in 1:Nstation){
        loglik.y[i] <- log(dnegbin(y[i], p[i], alpha))
      }
      }
      ")
sink()
inits<-function(){
  list(density = 5)
}
parameters<-c("density","rate", "alpha", "lambda","loglik.y", "loglik.stay")
n.chains<-3; n.iter<-2000; n.burnin<-1000; n.thin<-2
n.mcmc<-(n.iter-n.burnin)*n.chains/n.thin

model_neg<-jags(datalist, inits,parameters, model.file,
                n.chains=n.chains, n.iter=n.iter, n.burnin=n.burnin, n.thin=n.thin)
model_neg$summary[1:4,]

# Caluculate the WAIC value
waic(cbind(model_neg$sims.list$loglik.y, model_neg$sims.list$loglik.stay))

# Poisson-exponential model  with a density covariate
x <- as.numeric(scale(d2$X))
datalist<-list(y = y, area = area, effort = effort, is.censored = is.censored, stay = stay, cens = cens, Nstay = Nstay, Nstation = Nstation, act = act, x = x)
model.file<-"~/model1.txt"
sink(model.file)
cat("model
      {
      for(i in 1:Nstay){
        is.censored[i] ~ dinterval(stay[i], cens[i])
        stay[i] ~ dexp(rate)
      }
      for(i in 1:Nstation){
        y[i] ~ dpois(lambda[i])
        log(lambda[i]) <- log(area) + log(effort[i]) + log(density[i]) + log(rate) + log(act)
        density[i] <- exp(alpha + beta*x[i])
      }
      rate ~ dgamma(0.01, 0.01)
      alpha ~ dunif(-5, 5)
      beta ~ dunif(-5, 5) 
      # to calculate the WAIC value
      for(i in 1:Nstay){
        loglik.stay[i]<-log(ifelse(is.censored[i] == 1, 1 - pexp(stay[i], rate), dexp(stay[i], rate)))
      }
      for(i in 1:Nstation){
        loglik.y[i] <- log(dpois(y[i],lambda[i]))
      }
      
      }
      ")
sink()
inits<-function(){
  list(alpha = 0, beta = 0)
}
parameters <- c("density","rate", "lambda", "loglik.y", "loglik.stay")
n.chains <- 3; n.iter<-2000; n.burnin<-1000; n.thin<-2
n.mcmc <- (n.iter - n.burnin)*n.chains/n.thin

model_poisC <- jags(datalist, inits,parameters, model.file,
                    n.chains=n.chains, n.iter=n.iter, n.burnin=n.burnin, n.thin=n.thin)
model_poisC$summary

# Caluculate the WAIC value
waic(cbind(model_poisC$sims.list$loglik.y, model_pois$sims.list$loglik.stay))
```
